# Proteomic profiling of zinc homeostasis mechanisms in *Pseudomonas aeruginosa* through data-dependent and data-independent acquisition mass spectrometry

**DOI:** 10.1101/2025.01.13.632865

**Authors:** Annaliese C. S. Meyer, Matthew R. McIlvin, Paloma Lopez, Brian C. Searle, Mak A. Saito

## Abstract

Zinc is central to the function of many proteins, yet the mechanisms of zinc homeostasis and their interplay with other cellular systems remain underexplored. In this study, we employ data-dependent acquisition (DDA) and data-independent acquisition (DIA) mass spectrometry to investigate proteome changes in *Pseudomonas aeruginosa* under conditions of different zinc availability. Using these methods, we detected 2143 unique proteins, 1578 of which were identified by both DDA and DIA. We demonstrated that most of the previously described Zn homeostasis systems exhibit proteomic responses that follow similar trends to those seen in transcriptomics studies. However, some proteins that are considered instrumental in Zn homeostasis, notably those in Zn transporter ZnuABC, were not detected by our methods, although other proteins of other uptake systems were abundant. Furthermore, changes in abundance of multiple Zn-metalloproteins and Zn-independent homologs were clearly observable, with respective increases and decreases when Zn was provided, though the magnitude of these changes varied. Most of the Zn-metalloproteins observed were located in one of two Zur-regulated operons between PA5534 and PA5541. This study provides a view of Zn homeostasis mechanisms that is complementary to existing transcriptomics investigations: as gene transcripts are not strictly proportional to the actual distribution of proteins within a cell, analysis of the proteome offers another way to assess the relative use and importance of similar or ostensibly redundant systems in different conditions and can highlight shifts in metal prioritization between metalloproteins.

## 1. Introduction

Zn is an important micronutrient for many prokaryotes, second only to iron among d-block metal protein cofactors. In prokaryotes, Zn metalloproteins accounts for an average of 5-6% of all proteins for a given organism [1]. Correspondingly, it serves as the metal cofactor for a significant proportion of metalloproteins: for example, metalloproteomic analysis of the marine bacterium *Pseudoalteromons* revealed that nearly half of the identified metalloproteins were metalated with Zn [2]. Zn-binding proteins have a range of roles, from catalytic to regulatory.

Both the number of Zn-associated metalloproteins and the intracellular concentrations of Zn vary widely between species (reports range from <0.1 – 1 mM [3,4]). Maintaining tight control over Zn homeostasis is essential for proper cellular functioning. If there is insufficient Zn available to an organism, the proteins that require it for function cannot operate, and when free Zn in excess, it can prevent other transition metals from binding to their appropriate ligands by out-competing them for the position [5], thus preventing either cellular uptake of the metal or enzyme functionality (e.g. [6]). There are many and varied homeostatic mechanisms employed by prokaryotes [7–11] to exert this fine control on intracellular Zn concentrations, some of which are well characterized (e.g. the *znu* high-affinity ATP-binding cassette (ABC) transporter), and some of which have likely not yet been discovered. The loss of Zn-related uptake and efflux systems, while in some cases compensated by redundant systems, can exert a measurable growth defect [8,10].

*Pseudomonas aeruginosa* MPAO1 is a model prokaryote that contains several proposed and confirmed Zn uptake and efflux systems. Investigations of *P. aeruginosa* are most often contextualized in relation to human health concerns, as it is often implicated in nosocomial infections and is major cause of chronic infection in cystic fibrosis patients; however, *P. aeruginosa* is effectively ubiquitous in the environment [12]. Zn homeostasis in *P. aeruginosa* is primarily governed by the Zn uptake regulator (Zur, PA5499), which is a metalloregulatory protein in the ferric uptake regulator (Fur) protein family [13–16]. In Table 1, we summarize the known Zn efflux and uptake pathways and their component proteins. Of note in *P. aeruginosa* is the pseudopaline mechanism of Zn acquisition. Pseudopaline is an opine carboxylate Zn-chelating molecule, and is synthesized, released, and taken up by proteins within the *cnt* operon [17]. Recent studies have implicated pseudopaline as one of the primary pathways of Zn uptake in *P. aeruginosa* (e.g. [13]) and comparable molecules have been found in other prokaryotes, such as *Staphylococcus aureus* and *Yersinia pestis* [18]. The proteins in the *cnt* operon for pseudopaline synthesis and use are also conserved; homologs have been identified in approximately 250 species using genome neighbourhood network analysis [19].

**Table 1.** Proteins involved in Zn uptake and efflux in P. aeruginosa.

| Name | Function |
| --- | --- |
| *Uptake* |  |
| ZnuABC | High-affinity ABC transporter |
| ZnuD | TonB-dependent receptor |
| PA4063-4066 | Putative ABC transporter |
| CntI | Pseudopaline exporter |
| CntO | Pseudopaline importer |
| CntLM | Pseudopaline synthesis proteins |
| PA1922 | Putative TonB-dependent receptor |
| PA2911 | Putative TonB-dependent receptor |
| PA2912-2914 | Putative ABC transporter |
| HmtA | Atypical P-type ATPase importer |
| *Efflux* |  |
| CzcCBA | Efflux pump |
| CzcD | Cation diffusion facilitator transporter |
| YiiP | Cation diffusion facilitator transporter |
| CadA | P-type ATPase transporter |

In this study, we leverage two separate mass spectrometry approaches to interrogate the effects of variable Zn concentrations on the *P. aeruginosa* proteome. Liquid chromatography coupled to tandem mass spectrometry (LC-MS/MS) forms the basis of a large range of proteomic methods. One of the primary techniques for proteomics employs (liquid chromatography coupled to tandem mass spectrometry) LC-MS/MS instruments and applies a data-dependent acquisition (DDA) mode for peptide and protein analysis. For each cycle in a DDA run, precursor (unfragmented) peptide ions are measured in an MS1 survey and a number of the most abundant peptide ions are subsequently fragmented and measured to ideally produce one MS2 spectrum per peptide ion [20,21]. These spectra can be matched to peptide sequences and the count of the number MS2s assigned to a protein is used as a semi-quantitative measurement. A dynamically maintained “exclusion” list is typically employed to avoid over-sampling the same peptides but can result in a compressed dynamic range.

An alternative technique known as data-independent acquisition (DIA) has gained popularity since its introduction in 2012. The nuances of this method are reviewed elsewhere (e.g. [20,21]). Briefly, in each cycle an initial MS1 spectrum is followed by a series of MS2 spectra tiled to span the entire m/z range of the MS1 scan. Contrary to DDA, multiple peptide precursors are co-fragmented in each MS2 window such that even low-abundance peptides can be detected and quantified. However, the increase in fragmentation greatly increases the complexity of the resultant spectra, such that overall identification requires alternate informatic approaches.

Here we present an analysis of *P. aeruginosa* with these two proteomic methodologies to explore its response to varying Zn conditions, outline the response of several known mechanisms of Zn stress homeostasis, and identify several under-characterized systems that may play a significant role in modulating Zn use and intracellular concentrations.

## 2. Materials and methods

### 2.1 Strains and growth conditions

*P. aeruginosa* strain MPAO1 was provided from the Manoil Lab *P. aeruginosa* Mutant Library [22]. Cells were streaked from glycerol stocks on LB agar plates. To prevent zinc carry-over from LB plates into the experimental media and allow cells to acclimatize, single colonies were first transferred to standard M9 medium (M9 minimal medium per litre; 6 g anhydrous Na_2_HPO_4_ (Baker & Adamson), 3 g KH_2_PO_4_ (Fisher Scientific), 1 g NH_4_Cl (Sigma), 0.5 g NaCl (Fisher Scientific), trace salts solution as described in Harwood and Cutting [23]) lacking Zn and Co, then incubated overnight at 37 °C to an OD600 of 0.055 and 0.056 respectively. Spectrophotometric measurements were made using a SpectraMax 384 Plus spectrophotometer.

Then, 100 µL of the overnight cultures were added in triplicate to 250 mL new polypropylene Erlenmeyer flasks (US Plastic Corporation) that had been soaked in 1% Citranox, soaked in 10% trace metal clean HCl and rinsed three times with HCl diluted to pH 2. Each flask contained 15mL of M9 medium with or without 14.2 µM Zn(II) added. Cultures were grown at 37 °C with shaking at 140 RPM and had regular withdrawals of 200 µL for OD_600_ measurements, made using a 96-well plate. 4 mL of the cultures were harvested after 16 hours of growth while the cells were in mid-log phase using an Eppendorf 5810 R centrifuge at 180 x g for 20 minutes.

Growth curves are illustrated in supplemental Figure S1. Pellets were then frozen at -80 °C until further use.

### 2.2 Protein digestion

Protein extraction was performed based on the method described by Hughes et al. [24]. All reagents were prepared using Optima LC-MS grade water. Briefly, thawed cells were re-pelleted by centrifugation for 20 minutes at 14 100 x g in an ethanol-washed low adhesion 2 mL microcentrifuge tube (USA Scientific, part 1420-2600). 430 µL lysis buffer (50 mM HEPES pH 8.5 (Boston BioProducts)/1% SDS (Fisher Scientific)) was added to the pellet, mixed, and then 30 µL was withdrawn for protein quantification. Cells were then treated with 50 units of benzonase nuclease (Novagen) to degrade chromatin and incubated at 37°C for 40 minutes. The samples were incubated for 30 minutes at 45 °C with 20 µL of 200 mM dithiothreitol (DTT) (Fisher Bioreagents) in 50 mM HEPES pH 8.5 to reduce disulfide bonds, then for 30 minutes in the dark at 24°C with 40 µL of 400 mM iodoacetamide (Acros Organics) in 500 mM HEPES pH 8.5 to alkylate sulfhydryl groups. The iodoacetamide reaction was quenched with 40 µL of the DTT solution. Protein clean-up was then performed using a mixed magnetic bead solution of 0.70-1.10 µm and 1 µm average particle size (GE Healthcare UK Limited). 400 ug magnetic beads were added per 400 µL lysate. The solution was acidified to pH 2 using formic acid and then protein binding to the beads was induced with the addition of 1100 µL of 100% acetonitrile. The samples were then incubated at 37 °C in an Eppendorf Thermomixer R for 15 minutes, followed by 30 minutes at room temperature and 2 minutes on a magnetic rack. After removing the supernatant, samples were washed with 1400 µL of 70% ethanol followed by 1400 µL of 100% acetonitrile. The beads were then dried until the acetonitrile evaporated and reconstituted in 90 µL of 50 mM HEPES pH 8. The beads were incubated at room temperature for two hours. Proteins were quantified using a Micro BCA protein assay kit (Thermo Scientific) and a Nanodrop ND-1000 Spectrophotometer. A bovine serum albumin standard (Thermo Scientific) was used to create a standard curve (R^2^=0.997). Trypsin (Promega, V5280) was added in a ratio of 1:20 trypsin to protein, then samples were incubated at 37 °C for 20 hours. The digested samples were then washed using a final concentration of 95% acetonitrile then with 100% acetonitrile, then subsequently air-dried. The dried beads were reconstituted in 2% DMSO for a final protein concentration of 1.1 µg/µL. Following a 15-minute incubation of a magnetic rack, the supernatant was transferred to a clean tube, and then again transferred after a 5-minute incubation. 1% formic acid was then added to a final concentration of 0.1% and a 1 µg/µL protein concentration. Samples were then frozen at -20 °C until analysis.

### 2.3 Chromatography and Mass Spectrometry

Tryptic peptides were analyzed using liquid chromatography coupled with tandem mass spectrometry (LC/MS/MS) where a Michrom Advance HPLC reverse phase chromatography system was paired with a Thermo Scientific Q-Exactive Orbitrap mass spectrometer using a Michrom Advance CaptiveSpray source. Before injection, samples were concentrated onto a C18 trap column (0.2 x 10 mm ID, 5 µm particle size, 120 Å pore size, C18 Reprosil-Gold, Dr. Maisch GmbH), rinsed with 100 µL of a 0.1% formic acid, 2% acetonitrile (ACN) and 97.9% water solution before gradient elution with a reverse phase C18 column (0.1 x 250 mm ID, 3 µm particle size, 120 Å pore size, C18 Reprosil-Gold, Dr. Maisch GmbH) using a 500 nL/min flow rate. All solvents were prepared with Fisher Optima grade reagents. Liquid chromatography was a nonlinear 240-minute gradient from 5% to 95% buffer B, where A was 0.1% formic acid in water and B was 0.1% formic acid in ACN. Two modes of mass spectrometry were used to acquire data. During DDA analysis, MS1 scans proceeded from 380 m/z to 1280 m/z using a 70K resolution. The top 10 ions underwent MS2 scans with a 2.0 m/z isolation window and a 15-second dynamic exclusion time. During DIA analysis, the MS1 scan range was 385 m/z to 1015 m/z with a 60K resolution. The DIA scans comprised a precursor range of 400 m/z to 1000 m/z with 25 non-overlapping windows of 24 m/z. The MS2 resolution was 30K.

#### 2.3.1 Spectral processing and annotation

##### 2.3.1.1 DDA processing

The generated mass spectra from DDA analysis were searched against the *P. aeruginosa* PAO1 translated genome [25] using the SEQUEST HT algorithm within Proteome Discoverer 2.2 (Thermo Scientific). We used a fragment tolerance of 0.02 Da, a parent tolerance of 10.0 ppm, and a +57 carbamidomethyl fixed modification on cysteine. Variable modifications were a +16 oxidation modification on methionine and a +42 acetyl modification on the N-terminus. A maximum of two missed cleavages were allowed. Identification thresholds were 95.0% for peptides and 99.0% for proteins (minimum two peptides), giving a protein false discovery rate of 0.4% using decoys generated from a reverse database. Annotation and processing were performed using Scaffold 5.0.1 (Proteome Software, Portland, Oregon, USA). All data is presented as normalized total spectral counts.

##### 2.3.1.2 DIA processing

The spectra from DIA analysis were processed using several tools within ScaffoldDIA (Proteome Software, Portland, Oregon, USA). First, staggered windows were deconvoluted using ProteoWizard (3.0.11748) [26]. Spectra were searched using the FASTA searching module against the *P. aeruginosa* PAO1 translated genome [25] using a 10.0 ppm peptide mass tolerance and a 10.0 ppm fragment mass tolerance, and only peptides of length 6 to 30 and charge 2 to 3 were included. A maximum of 1 missed cleavage was permitted.

False discovery rates were calculated using decoys generated from a reverse database.

Percolator [27–29] (3.01) was used to filter peptides in each sample to achieve a maximum false discovery rate of 0.01, then individual search results were combined, and posterior error probabilities were generated for all peptide spectral matches. These results were filtered to a false discovery rate threshold by Percolator. EncylopeDIA [30] (1.12.31) was used to quantify peptides using the 5 fragment ions of the highest quality. Protein groups were created if peptides corresponded well to more than one protein. Protein identifications were filtered to a threshold false discovery rate of <1.0%. Normalized exclusive intensity values were used for further analysis.

#### 2.3.2 Statistical Analysis and Visualization

To compare the fold-change values as determined by both DIA and DDA, we used a pseudocount of 0.5 for proteins in DDA where one condition had zero spectral counts. We then performed a paired two-tailed t-test on the mean fold-change between treatments, which indicated that there was no significant global difference between values determined by both methods (p-value > 0.1). Differences that do exist between these methods are likely due to a combination of the compressed dynamic range of DDA and this artificial increase in spectral counts, as the calculated fold-change will change depending on the chosen denominator.

Functional enrichment and protein network scores were computed using STRING v12.0 [31–37] and calculated based on the set of log_2_ fold-change (LFC) values of all proteins detected using DIA methods. Initial p-values for each functional grouping were determined by the Kolmogorov-Smirnov test or aggregate fold change test. FDR was determined using the Benjamin-Hochberg procedure. Only groupings with an FDR < 0.05 were reported. Full results for both DIA and DDA enrichment analyses are available as supplemental files.

We performed a heteroscedastic two-tailed t-test to assign p-values to the comparison of Zn-replete and Zn-depleted samples for each protein using a significance level of α = 0.1. At this α, the minimum significant effect size is 2.5 (Cohen’s d) at a power of 0.8 [38]. We identified proteins of interest based on those proteins within enriched groups as determined by the STRING analyses (Table S1), which led to the investigation of known and suspected Zn trafficking proteins, Zn-metalloproteins, cell envelope modification systems, and proteins in pathways or gene clusters associated with those identified by the aforementioned methods. In Table 2, we summarize the individual results for proteins of interest. The raw data is available through the ProteomeXchange Consortium via the PRIDE partner repository with the dataset identifier PXD047454 and 10.6019/PXD047454. Processed results are available in a supplemental file.

**Table 2.** Relative abundances of investigated proteins between Zn-depleted and Zn-replete conditions (LFC = log_2_(Zn-depleted/Zn-replete)) and associated statistical measures. P-values were determined through a heteroscedastic two-tailed t-test. Bolded values exceed desired statistical thresholds of p-value < 0.1 and effect size > 2.5. Missing values indicate the protein in question was not detected by that methodology.

| Locus Tag | Protein Name | DDA |  |  | DIA |  |  |
| --- | --- | --- | --- | --- | --- | --- | --- |
|  |  | LFC | p-value | Cohen's *d* | LFC | p-value | Cohen's *d* |
| PA0781 | ZnuD | 9.477 | **0.031** | **4.528** | 6.692 | **0.04** | **3.964** |
| PA0958 | OprD | 0.984 | **0.072** | 1.995 | 0.743 | **0.063** | 2.107 |
| PA1922 | - | 5.341 | **0.038** | **3.961** | 4.141 | 0.128 | 2.048 |
| PA1923 | - | 3.92 | 0.287 | 1.096 | 3.852 | **0.076** | **2.757** |
| PA1925 | - | - | - | - | 4.172 | 0.149 | 1.856 |
| PA2435 | - | 2.87 | **0.052** | **2.955** | 2.903 | 0.121 | 2.098 |
| PA2437 | - | 1.712 | 0.423 | 0.567 | - | - | - |
| PA2438 | HflC | 4.945 | **0.011** | **6.284** | 3.601 | **0.053** | **3.366** |
| PA2439 | HflK | 2.734 | 0.184 | 1.387 | 3.788 | **0.065** | **3.015** |
| PA2505 | OpdT | -3.927 | **0.055** | **2.984** | -2.787 | 0.181 | 1.639 |
| PA2520 | CzcA | -4.406 | 0.13 | 1.94 | -1.924 | 0.103 | 2.226 |
| PA2521 | CzcB | -4.396 | 0.203 | 1.452 | -3.476 | 0.112 | 2.221 |
| PA2522 | CzcC | -2.583 | **0.041** | **3.268** | - | - | - |
| PA2523 | CzcR | -1.446 | 0.423 | 0.517 | - | - | - |
| PA2524 | CzcS | - | - | - | 0.045 | 0.939 | 0.07 |
| PA2911 | - | 5.212 | 0.12 | 2.081 | 5.728 | **0.094** | 2.475 |
| PA2913 | - | 2.855 | 0.309 | 0.95 | 6.686 | **0.077** | **2.767** |
| PA3438 | FolE1 | 0.752 | 0.748 | 0.285 | - | - | - |
| PA3527 | PyrC1 | -1.69 | **0.039** | **3.063** | -0.396 | 0.56 | 0.52 |
| PA3574.1 | CopZ2 | -1.255 | **0.054** | 2.207 | -1.299 | 0.225 | 1.277 |
| PA3601 | RpmE2 | 1.39 | 0.423 | 0.505 | 4.841 | **0.044** | **3.714** |
| PA3690 | CadA/ZntA | -6.113 | **0.013** | **7.029** | -3.332 | **0.002** | **11.541** |
| PA4022 | HdhA | -7.787 | **0.015** | **6.488** | -2.000 | **0.064** | 2.127 |
| PA4063 | - | 7.291 | **0.017** | **6.233** | 5.857 | **0.019** | **5.856** |
| PA4064 | - | 4.302 | 0.361 | 0.911 | - | - | - |
| PA4065 | - | 4.87 | **0.068** | **2.873** | 4.305 | **0.005** | **11.119** |
| PA4066 | - | 1.39 | 0.423 | 0.505 | 3.202 | 0.132 | 1.993 |
| PA4381 | ColR | -3.172 | **0.07** | **2.596** | -4.548 | 0.362 | 0.958 |
| PA4723 | DksA | -1.356 | **0.055** | **3.07** | -0.03 | 0.969 | 0.035 |
| PA4837 | CntO/ZrmA | 5.981 | **0.017** | **6.098** | 4.642 | **0.018** | **5.666** |
| PA5049 | RpmE | -1.156 | **0.068** | 2.164 | -2.966 | **0.093** | 2.462 |
| PA5105 | HutC | -0.403 | 0.423 | 0.199 | -0.581 | 0.406 | 0.759 |
| PA5499 | Zur | -3.503 | **0.066** | **2.757** | -0.181 | 0.703 | 0.344 |
| PA5531 | - | 3.22 | **0.004** | **11.162** | 1.077 | 0.183 | 1.526 |
| PA5534 | - | 3.478 | 0.423 | 0.743 | 4.255 | **0.056** | **3.309** |
| PA5535 | ZigA | 6.304 | **0.045** | **3.662** | 5.838 | **0.084** | **2.639** |
| PA5536 | DksA2 | 1.712 | 0.423 | 0.567 | 0.867 | 0.413 | 0.746 |
| PA5539 | FolE2 | 3.214 | 0.212 | 1.318 | - | - | - |
| PA5540 | - | 3.802 | **0.077** | **2.562** | - | - | - |
| PA5541 | PyrC2 | 0.434 | 0.423 | 0.212 | - | - | - |

The Pathway Tools software (v.28) and BioCyc databases were used to generate the genome overview figures and summary metabolic maps, based on the *P. aeruginosa* PAO1 gcf_000006765cyc v.29 database (NCBI Accession: PRJNA57945) [25,39–55].

## 3. Results and Discussion

### 3.1 Global proteomic response

We identified 1883 proteins using DDA methods and 1838 proteins using DIA methods. 1578 proteins, or 74%, were identified by both methods, while 14% and 12% of proteins were unique to DDA and DIA methods, respectively. Unique identifications are likely attributable to the stochastic identification of rare proteins. The high proportion of shared proteins suggests that both methodologies provided an accurate and comparable view of the proteomic response of *P. aeruginosa* to Zn. By using both methodologies, we take advantage of the strengths of each while compensating on their relative weaknesses. The robustness of single precursor identification available through DDA and the sensitivity to low abundance peptides offered by DIA – leading more precise fold-change values - makes these methods complementary.

Additionally, using two independent analyses provide more support to trends observed through both methods. As such, fold-change values reported in this manuscript are calculated from DIA-derived exclusive intensities where available, and relevant proteins identified by DDA but not detected by DIA are discussed using DDA spectral counts.

We first assess the response of the entire *P. aeruginosa* proteome to Zn-depleted and Zn-replete treatments. The proteins identified as significantly different between treatments are not limited to only those directly involved in Zn trafficking or use: we observed notable changes across several core systems and pathways. This high-level approach allows us to place the changes we measured in known or suspected Zn homeostasis mechanisms within the larger context of cellular stress responses and metabolic processes induced or affected by Zn concentration. We then focus our remaining discussion on some of the major themes indicated by the functional enrichment analyses (Section 3.1.1), that is, transport of Zn and other metal ions, Zn-binding proteins and their homologs, and cell envelope maintenance and modification. The log_2_ fold-change of protein abundances are depicted as an overlay of the *P. aeruginosa* genome (Figure 1, DDA) such that the co-location of genes for which the protein products were differentially abundant can be visualized. Summary metabolic maps, available as supplemental files, illustrate the relative increase or decrease of individual proteins in major pathways.

**Figure 1.**
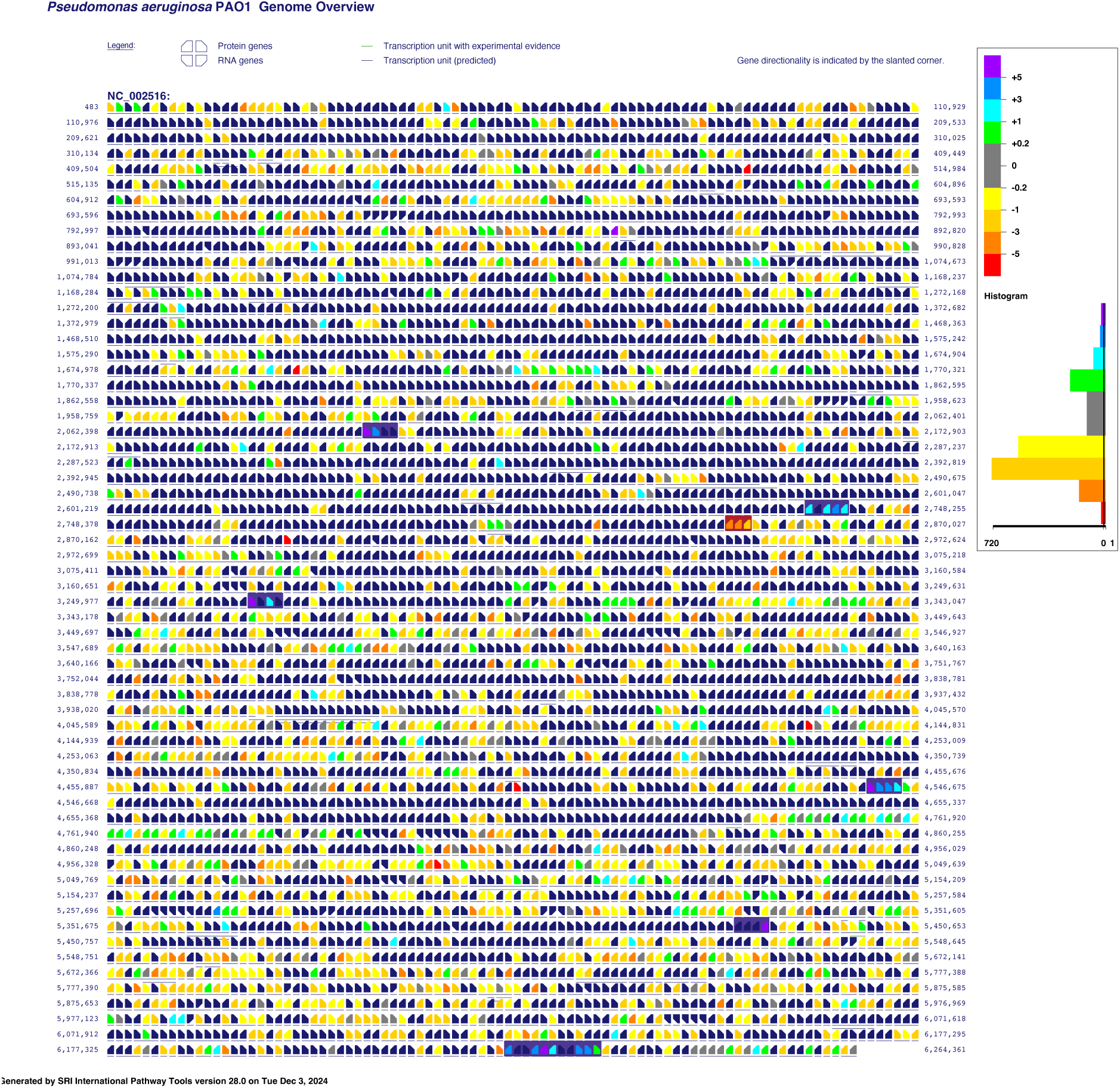
Genome overview of P. aeruginosa PAO1 overlaid with log_2_ fold-change abundance values as determined through data-dependent acquisition. Positive log_2_ fold-change values (cool colours) indicate enrichment in the Zn-depleted samples, and negative fold-change values (warm colours) indicate enrichment in the Zn-replete samples. Notable operons discussed in this article are highlighted in dark purple for those involved in Zn-acquisition and Zn-sparing functions, and in burgundy for those involved in Zn-efflux and Zn-resistance processes. Direction of transcriptions is indicated by the slant of gene icon. Figure was generated using SRI Pathway Tools v.28 and BioCyc database P. aeruginosa PAO1 gcf_000006765cyc v.29.

#### 3.1.1 Perturbation of cellular systems and functional enrichment across treatments

Detected proteins were associated with overarching cellular systems and functional groupings, as described by Gene Ontology (GO) Biological Process and Molecular Function terms, as well as STRING [31–37] (citation) local network clusters (Figure 2). Under Zn-replete conditions, there was broad, low-level enrichment of translation, protein export, and related amide metabolic processes, driven in part by the large number of ribosomal proteins that were more abundant in these samples. Proteins associated with ATP synthesis were also enriched.

**Figure 2.**
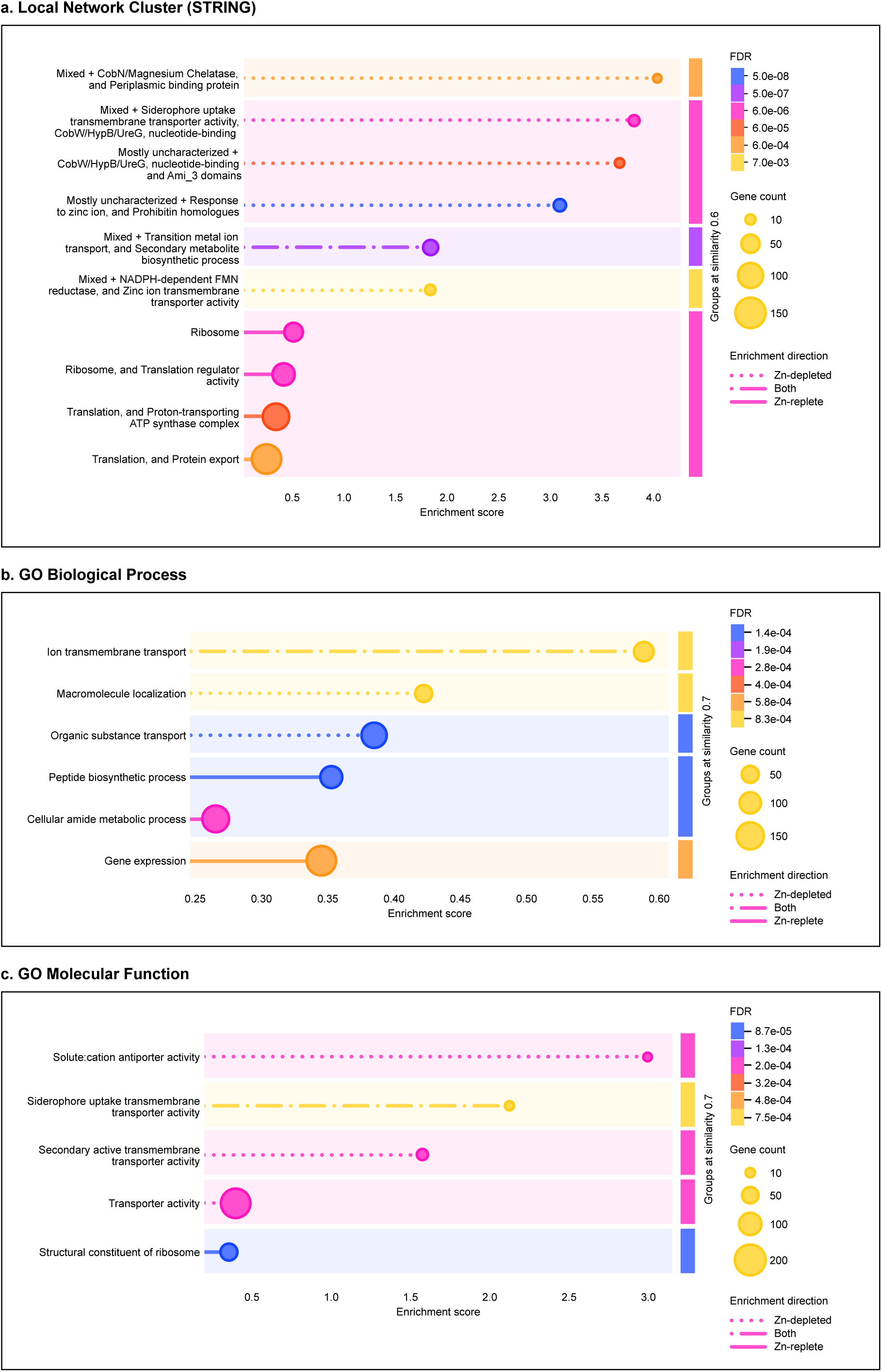
Perturbed functions and processes between Zn-depleted and Zn-replete treatments of *P. aeruginosa*. Function or process descriptions were merged if similarity exceeded 0.8. Terms were grouped at a similarity level of 0.6 (a) or 0.7 (b,c). Dotted lines indicate enrichment in the Zn-depleted condition, solid lines indicate enrichments in the Zn-replete condition, and combination dash-dot lines indicate enrichment in both treatments. The length of each line corresponds to a group’s enrichment score, or the degree to which a group is perturbed over the expected background. Colours represent the false discovery rate associated with the enriched group as determined by the Benjamin-Hochberg procedure. Marker size is scaled to the number of proteins detected that are associated with the enriched group. Panel (a) depicts enriched local network clusters as defined by STRING v12.0, panel (b) depicts enriched Gene Ontology Biological Process terms, and panel (c) depicts enriched Gene Ontology Molecular Function terms.

While there was some enrichment of siderophore and ion transmembrane transport systems in the Zn-replete samples, the enriched terms in the Zn-depleted samples were overwhelmingly associated with transporter activity, particularly metal ion transporter activity, and with Zn response; further, they generally exhibited higher enrichment scores as compared to the scores of enriched systems in the Zn-replete treatment. It is also worth noting that many proteins experimentally determined to be associated with Zn trafficking are not broadly annotated as such, so may not be well-represented by GO terms and should be considered based on STRING networks and experimental evidence.

Nevertheless, most of the individual proteins with large changes in abundance between treatments were those that were relatively higher in the Zn-replete samples: for example, through DIA measurements, 105 proteins were upregulated in the Zn-replete samples with an effect size greater than the detectable limit of 2.5, whereas only 48 proteins with comparative increases under Zn-depleted conditions exceeded that threshold. This suggests that the stress due to Zn starvation caused cellular resource allocation to shift towards Zn-acquisition and Zn-sparing functions, while a wider variety of cellular processes could be undertaken under Zn-replete conditions. That the Zn-depletion imposed by these experimental conditions was sufficient to cause such stress is supported by the small discrepancy in growth rates between treatment groups (Figure S1). In the Zn-replete condition, the growth rate was 7.71 ± 0.59 day^-1^ (n=3). In the Zn-depleted condition, the growth was slower, at 6.17 ± 1.01 day^-1^ (n=3). Both rates were calculated during exponential phase.

### 3.2 Zn acquisition

There are numerous systems for Zn uptake in *P. aeruginosa*, outlined briefly in Table 1.

This section describes some of these systems that were observed to be differentially abundant between the Zn-depleted and Zn-replete conditions (Figure 3). Of the eight systems of Zn uptake described in recent analyses of Zn homeostasis [15,56,57], components of seven were detectable in this study. Briefly, all four of the TonB-dependent receptors previously implicated in Zn import were detected and followed expected patterns of enrichment in Zn-depleted samples [15]. Out of three expected ABC transporter systems, only the ZnuABC system remained elusive.

**Figure 3.**
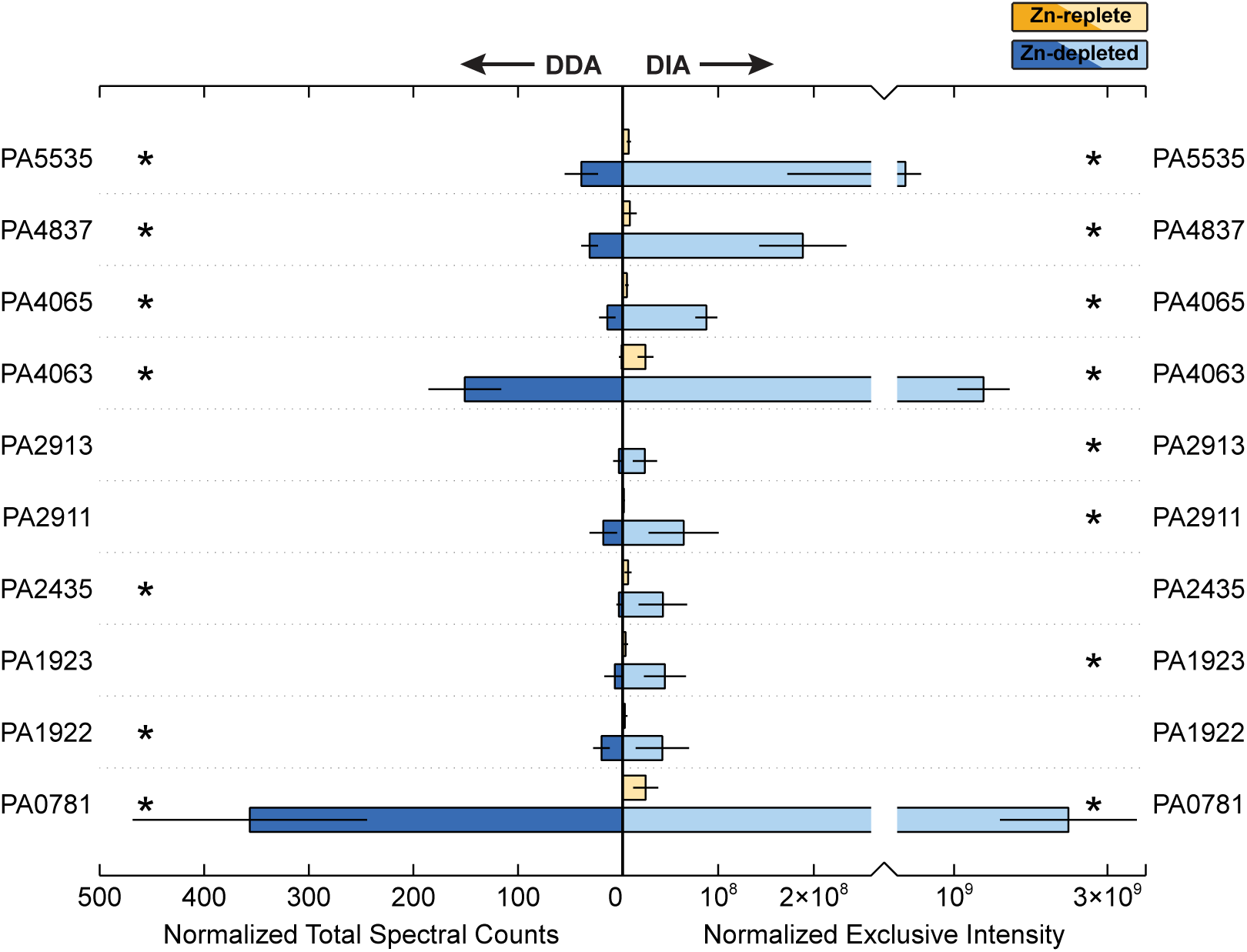
Relative abundance of proteins involved in Zn acquisition between the Zn-depleted and Zn-replete treatments, averaged across biological triplicate samples. Normalized total spectral counts as determined through data-dependent acquisition are on the left, and normalized exclusive intensity as determined through data-independent acquisition is on the right. Yellow shades correspond to the Zn-replete samples and blue shades to the Zn-depleted samples. Error bars represent one standard deviation calculated from the triplicate samples. Stars indicate proteins for which the difference in abundance between the two treatments is statistically significant (p-value < 0.1).

ZnuABC is the most common Zn acquisition in bacteria, and broadly distributed [7,8,10,15,16,58]; though undoubtedly an important Zn uptake strategy in *P. aeruginosa*[15,16,56,58], its absence here suggests that under these conditions, this conserved system is less advantageous than its other transporters. The final expected system, P-type ATPase HmtA, was also detected, and will be discussed later in this manuscript. We also report a possible Zn metallochaperone, PA5535.

The abundance of PA0781, or ZnuD [16,59] was elevated in the Zn-depleted conditions (Figure 3). There was an approximately 100-fold increase in DIA signal in the Zn-depleted samples; as one of the most differentially abundant proteins observed, PA0781 may serve as primary mode of entry for extracellular Zn. Our results corroborate those of earlier studies which found that transcripts of *znuD* increased 172-fold when the *znuA* gene was knocked out, which limited intracellular Zn concentrations by approximately 60% in *P. aeruginosa* [16].

A second TonB-dependent receptor, PA1922, was also more abundant in the Zn-depleted conditions, increasing 18-fold against the Zn-replete condition. Pederick et al. [16] also observed upregulation of this gene in their Δ*znuA* mutant. PA1922 may serve as a redundant system alongside ZnuD for import of free Zn(II) ions [56], We also detected PA1923 and PA1925, which, alongside the here-undetected protein PA1924, likely form an associated import system [55]. Similarly, these proteins were abundant only in the Zn-depleted condition. Intriguingly, PA1923 contains a CobN-like or cobaltochelatase-like domain. As the PA1922-1925 Zn import system lacks orthologs within most other members of the *Pseudomonas* genus [56], and given that Co(II) and Zn(II) share some similar chemical characteristics [60–63], we speculate that perhaps the PA1922 derived from a Co(II) import system that has been sacrificed in *P. aeruginosa* to form an additional import system for Zn acquisition, and/or that developed promiscuity in metal selection, allowing it to transport both metals.

The CntO TonB-dependent receptor (PA4837 or ZrmA, [13,64]) permits the import of pseudopaline complexed with Zn [13]. CntO was enriched 25-fold in Zn-depleted conditions (Figure 3). A similar pattern was observed in a recent transcriptome study, where CntO showed a 110-fold increase under Δ*znuA*-induced Zn depletion [16]. Interestingly, CntO transcripts were also upregulated 8.95-fold in response to human airway mucus secretions and was also induced in sputum of cystic fibrosis patients [65,66]. These secretions are known to contain calprotectin, produced by neutrophils [67], which can sequester Zn and thus present a Zn-deficient environment to colonizing bacteria [66]. Despite the strong response of CntO to the Zn-depleted treatment, the other proteins in this operon were not detectable. These proteins are responsible for the synthesis and export of pseudopaline. Future studies using more sensitive two-dimensional LC-MS and targeted proteomic methods may be able to detect these proteins under similar conditions.

A final TonB-dependent receptor, PA2911, increased 53-fold in Zn-depleted samples. PA2911 is co-located in the *P. aeruginosa* genome with the PA2912-PA2914 ABC transporter. Of that transporter system, only PA2913, a likely solute binding protein [16,59,68], was detectable and notably enriched. Recently, Secli and colleagues suggested that the PA2911-2914 system could be responsible for the import of pyochelin-bound metal. Pyochelin is capable of binding multiple divalent cations, including Zn^2+^ and Co^2+^ [69,70], though it is primarily considered in the context of Fe^2+^ import. The canonical pyochelin importer, FptA, imports pyochelin bound to metals other than Fe only at extremely low efficiencies [68], so the existence of a Zur-regulated system for the import of pyochelin bound to Zn and/or Co seems plausible.

The PA4063-4066 cluster encodes a putative Zn ABC-transporter, several components of which were more abundant in the Zn-depleted condition (Figure 3). In the case of PA4063 and PA4065, DIA exclusive intensity increased by approximately 58-fold and 20-fold respectively against the Zn-replete treatments. The designation of the PA4063-4066 gene cluster as a Zn-importing ABC transport system is supported by observations of its upregulation in response to Zn limitation in a transcriptome study and the presence of a Zur (PA5499) binding site (e.g. [15,16,56]). Zur is observed in both treatments through DIA, and present in the Zn-replete samples in DDA, though with extremely low signal.

PA4063 has been tentatively suggested as a solute binding protein (SBP) for this system, a periplasmic Zn chaperone, or metal sensor [15,16,71] in part due to the presence of histidine-rich loops that gain structure in the presence of Zn^2+^, binding two ions with relatively low affinity. PA4066 is another possible SBP candidate [15,16,56]. Though PA4066 was modestly enriched in the Zn-depleted samples, there was minimal signal through both the DDA and DIA methods. We suggest that the comparatively weak response of PA4066 to Zn depletion is evidence against its assignment as an SBP. PA4064 was similarly weakly increased in the Zn-depleted condition. As PA4064 has been annotated as the ATP-binding portion of the transporter and PA4065 as the permease [59], if this cluster is indeed an Zn ABC transporter, the function of PA4066 remains obscure.

Alternatively, Secli and colleagues [68] recently suggested that the PA4063-4066 cluster could encode for a MacB transporter responsible for the export of pyochelin. The accumulation of free metallophores in the cycloplasm and/or periplasm is disadvantageous under conditions of metal starvation, as it both prevents those metallophores from being able to bind metals for import extracellularly and allows them chelate metals within the cell, sequestering them from use. Secli et al. [68] note that pyochelin is crucial for cellular acquisition of Co, and further, that PA2914 and PA4065 are essential for maintaining intracellular Co homeostasis. However, adding deletions of PA2914 and/or PA4065 to *P. aeruginosa* strains where *znuA* and *cntL* were knocked out did not lead to a greater decrease in the intracellular Zn concentration, does lead to an increased cellular Zn demand [68]: they thus suggest that the PA2911-2914 and PA4063-4066 systems are primarily involved in the trafficking of pyochelin-Co complexes, and that the increase in these systems under Zn depletion reflect a greater need for Co, as it can in some cases function as a substitute for Zn. As Co was not provided in the media used in these experiments, that these systems were favoured over the ZnuABC transporter could provide some support towards their possession of dual function of Zn and Co acquisition. Further work in which proteomic studies of relevant *P. aeruginosa* mutants are matched with protein-metal allocations (‘metalloproteomics’, e.g. [2]) could reveal offer valuable insight into the function of these systems.

Metal-transporting P-type ATPase HmtA (PA2435) was enhanced under Zn depletion, in contrast to the unchanging mRNA values reported between various Zn concentration treatments in a recent transcriptomics study [15]. HmtA sits near a cluster of genes (PA2437-2439, PA2436 was not detected) where the corresponding protein abundances were also elevated in Zn-depleted samples, and significantly so for PA2438 (DDA) and PA2439. Though not robustly annotated, these appear to be HflC and HflK proteins respectively [16], which are known interactors with Zn-metalloprotease HflB [72] and are proximal to a potential Zur binding site [16]. We emphasize that the sensitivity and robustness of untargeted proteomic analyses, especially with paired methodologies, can complement transcriptomics data through detection of changes that may not be reflected in the mRNA signal.

The COG0523-domain containing gene, PA5535 (Cluster 1 [73]), is occasionally noted as differentially abundant in transcriptomic studies of Zn limitation or excess (e.g. [16]) and as with other cluster 1 COG0523 proteins, appears to be regulated by Zur [56,73]. However, due to an early association of all COG0523 proteins with CobW-like features, PA5535 is often represented as a cobalamin biosynthesis protein; however, based on sequence alignment, PA5535 is likely Zn metallochaperone GTPase ZigA (100% amino acid identity with other *Pseudomonas* sp. ZigA proteins), which is line with the observed dominance of ZigA and ZigA-like proteins in the cluster 1 COG0325 group [73]. In this work, PA5535 was abundant only in Zn-depleted media, with a 57-fold increase against the Zn-replete condition. Zn metallochaperones may assist in ensuring appropriate transfer of Zn to proteins with an absolute requirement for this metal under low Zn conditions. In *Acinetobacter baumannii*, ZigA interfaces with a histidine catabolic system, HutUHTIG [59,73,74] potentially acting to provide Zn to a histidine ammonia lyase, HutH. Capdevila et al. [3] noted that the HutUHTIG system is also present in *P. aeruginosa*, suggesting that ZigA-Hut mediated transport may be the reason for the observed upregulation of PA5535 in our study, and indicating a role for zinc in regulating the cellular glutamate pool. We observed the presence of the HutC (PA5105) transcriptional repressor in both DIA and DDA datasets, though at low intensities/spectral counts respectively. Previous studies similarly report depression of Hut operon proteins in response to divalent metals [75], including Zn in *P. putida* [76].

### 3.3 Zn Efflux and Resistance Strategies

As anticipated, proteins associated with Zn efflux and metal resistance were comparatively more abundant under Zn-replete conditions (Figure 4). While most of the commonly discussed Zn efflux systems were detected in this study, neither CzcD or Yiip, the cation diffusion transporters normally associated with Zn export in *P. aeruginosa*, were present in either treatment. However, all components of the well-characterized CzcCBA resistance-nodulation-division efflux pump system were enriched in the Zn-replete treatment, though CzcC (PA2522) was only detectable in the DDA data. CzcA (PA2520) and CzcB (PA2521) were 4-fold and 11-fold greater relative to the Zn-depleted samples, though not significantly, while the increased abundance of CzcC was significant. Previous works have also highlighted the importance of metal-transporting P-type ATPase CadA (PA3690, or ZntA) in Zn efflux, especially in early phases of Zn exposure, and identified an interplay between CadA and the CzcCBA system [15]. Fittingly, abundances of CadA increased 10-fold with the Zn-replete treatment.

**Figure 4.**
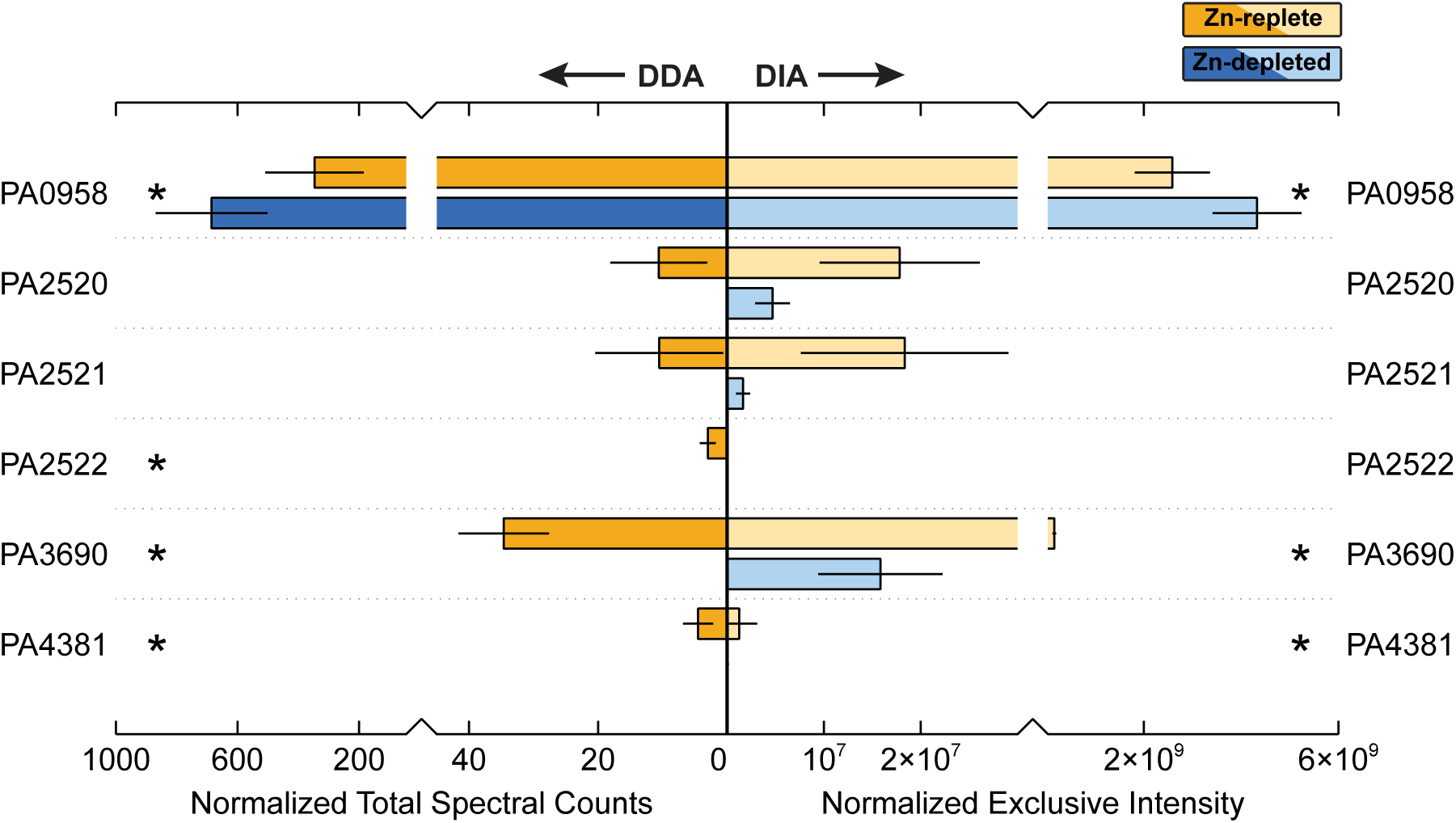
Relative abundance of proteins involved in Zn efflux and tolerance between the Zn-depleted and Zn-replete treatments, averaged across biological triplicate samples. Normalized total spectral counts as determined through data-dependent acquisition are on the left, and normalized exclusive intensity as determined through data-independent acquisition is on the right. Yellow shades correspond to the Zn-replete samples and blue shades to the Zn-depleted samples. Error bars represent one standard deviation calculated from the triplicate samples. Stars indicate proteins for which the difference in abundance between the two treatments is statistically significant (p-value < 0.1).

CzcRS, the two-component system (TCS) involved in the regulation of the CzcCBA pump, was observed in these data, though both components were exhibited very low signal and minimal change between treatments. However, OprD (PA0958), known to be repressed by CzcRS upon Zn exposure [77,78] did undergo a remarkable decrease in the Zn-replete condition. OprD allows for the uptake of positively charged amino acids and some antibiotics. While tempting to suggest that OprD may provide a route for incidental Zn uptake that must be closed when Zn is in excess, Zn exposure in other *Pseudomonas* species results in an increase in OprD expression [56,79]. However, *P. aeruginosa* appears to be somewhat unique within its genus in that Zur binding sites exists just upstream of the CzcR gene, which supports a potential role of OprD in Zn homeostasis.

### 3.4 Zn-sparing strategies in P. aeruginosa

While increasing the abundance of Zn uptake systems is clearly critical for cell survival in Zn-depleted environments, *P. aeruginosa* is further equipped with strategies to reduce its dependence on Zn overall. Numerous Zn metalloproteins in the *P. aeruginosa* genome are matched with homologous proteins that can either employ a different metal to achieve the same function, or do not require a metal cofactor at all. Within this study, we observe this strategy in action: there were marked differences in the abundance of some Zn-containing proteins between Zn-replete and Zn-depleted conditions, and conversely, the associated Zn-sparing homolog or analog was often detectable in the Zn-depleted samples. Figure 5 illustrates the abundance as measured via DDA of several Zn-dependent proteins and their Zn-independent partners between the two studied conditions.

**Figure 5.**
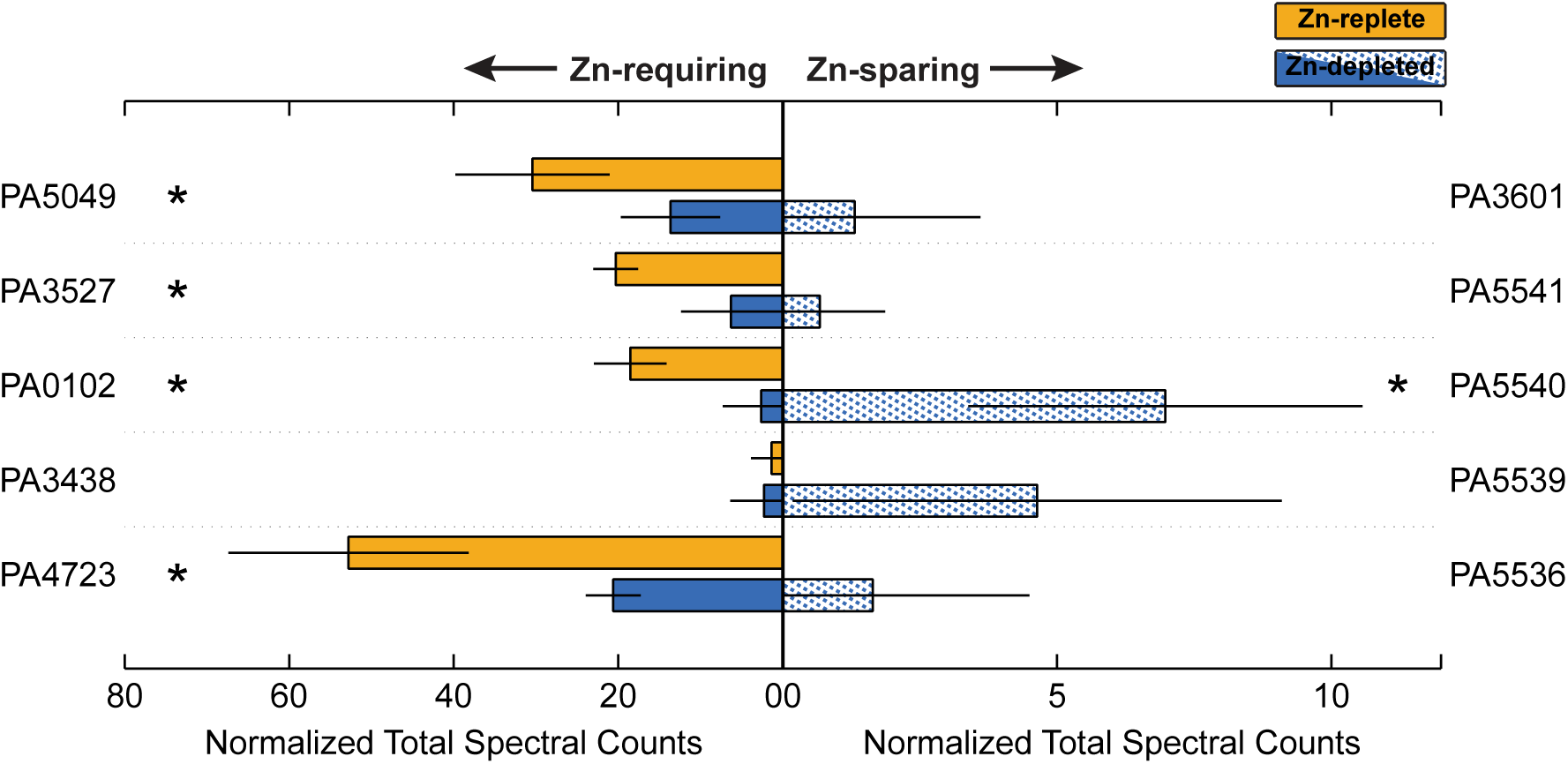
Relative abundance of selected proteins that require Zn (left) and the Zn-independent homolog or functional analog (right) between the Zn-depleted and Zn-replete treatments, averaged across biological triplicate samples. Data-dependent acquisition measurements are presented in the form of normalized total spectral counts as determined through data-dependent acquisition are on the left. Yellow shades correspond to the Zn-replete samples and blue shades to the Zn-depleted samples. Error bars represent one standard deviation calculated from the triplicate samples. Stars indicate proteins for which the difference in abundance between the two treatments is statistically significant (p-value < 0.1).

PA5535, the COG0523 Zn-metallochaperone discussed in the section 3.2, sits within a neighbourhood of other genes that also exhibit higher abundances in the Zn-depleted condition (Figure 6). Zur binding sites occur between PA5536 and PA5537, and between PA5538 and PA5539. PA5531, PA5534, PA5536, PA5539, PA5540 and PA5541 all display various degrees of enrichment (Table 2). PA5531 will be discussed further in section 3.5. PA5536, PA5539, PA5540, and PA5541 encode either metal-promiscuous or metal-independent paralogs or functional analogs of known Zn-metalloproteins. DksA2 (PA5536) is a paralog of DksA (PA4723), involved in the stringent response; FolE2 (PA5539) is a paralog of FolE1 (PA3438), a GTP cyclohydrolase; PA5540 is a putative γ-carbonic anhydrase, which could substitute for Zn-dependent β-carbonic anhydrases; and PyrC2 (PA5541), is a dihydrootase homologous to PyrC1 (PA3527) [15]. The Zn-dependent counterparts to these proteins are generally lower in abundance in Zn-depleted samples (Figure 5), with the exception of FolE1, for which there was no large change. Only one of the annotated β-carbonic anhydrases (PA0102) was markedly lower under Zn-depletion. PA5534 is not annotated but may encode another Zn-sparing paralog or Zn-chaperone. This Zn-sparing behaviour has been observed in other works, for example, Pederick et al. [16] also noted an upregulation of PA5540 under Zn-depleted conditions. Similar patterns were also observed in the homologous gene cluster in *Pseudomonas protegens* Pf-5 [80].

**Figure 6.**
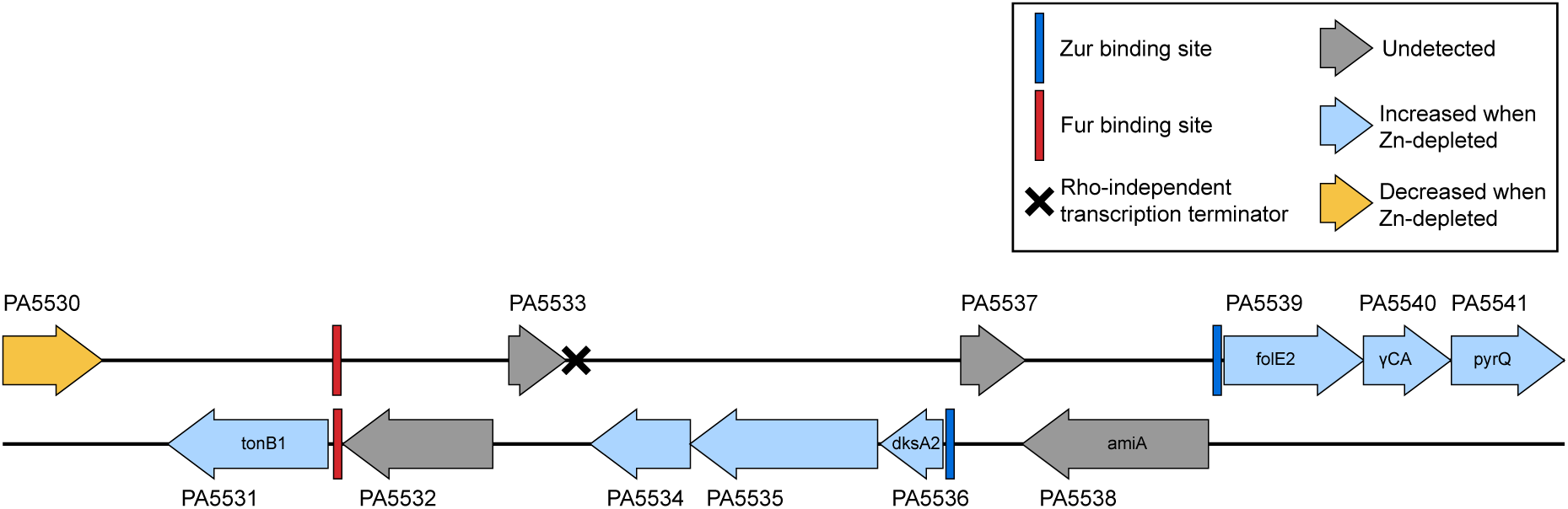
Several proteins encoded in the *P. aeruginosa* PAO1 genome gene cluster from PA5531 to PA5541 were more abundant in Zn-depleted conditions as compared to Zn-replete conditions. A Zur binding site (blue rectangle) is located on each strand upstream of the upregulated proteins. A Fur binding site (red rectangle) is immediately upstream of PA5531, the tonB1 gene. Gene annotations, the Fur binding site and the Rho-independent transcription terminator were assigned as indicated in the Pseudomonas Genome Database [26,90] for P. aeruginosa PAO1. The Zur binding sites were assigned by querying the genome for the Zur binding site sequences as determined for P. aeruginosa PAO1 by Pederick and colleagues [16].

As shown in in the functional enrichment and pathway perturbation analyses (Section 3.1.1), ribosomal proteins are broadly affected by Zn concentrations. For example, RpmE (PA5049), a Zn-containing L31 ribosomal protein [59], was much more abundant in the Zn-replete condition. RpmE function can be replaced by Zn-independent RpmE2 (PA3601) [16,81]. As expected, Zur-controlled RpmE2 is nearly undetectable in the Zn-replete treatment but present in the Zn-depleted treatment, consistent with the replacement of its Zn-requiring counterpart. If we consider the collective pool of RpmE and RpmE2 as derived from DIA exclusive intensities, when Zn is available, RpmE accounts for 99.8% of it. In contrast, 33.6% of the pool in the Zn-depleted treatment is Zn-independent RpmE2. This change is particularly dramatic, likely because RpmE2 exhibits comparable activity to RpmE (as shown in *E. coli* [82]): RpmE can be downregulated and its Zn reallocated without large deleterious effects. As a counterexample, the proportion of Zn-sparing DksA2 in the joint DksA/DksA2 pool increased only minimally, from 0.7% to 1.3%, between Zn-replete and Zn-depleted treatments. DksA regulates hundreds of genes, and though some reports shows that DksA2 can fully compensate for DksA in regulating the majority of its target genes in *P. aeruginosa* [83], others document that DksA2 has a higher comparative sensitivity to reactive oxygen species [84]. Further investigation of the regulation of these paralogs is beyond the scope of this work, but we suggest that as a constitutively expressed master regulator, DksA might nevertheless be prioritized as a Zn recipient, especially in the low nutrient conditions imposed by M9 minimal media. Moreover, DksA may be a relatively small reservoir of Zn compared to Zn-containing ribosomal proteins, thus limiting the utility of liberating DksA-bound Zn. Conversely, L31 and L33 proteins have been repeatedly described as Zn storage mechanisms [85–88], thus exchanging these proteins for Zn-independent paralogs provides a more substantial Zn supply.

### 3.5 Zn homeostasis mechanisms in non-canonically Zn-associated systems

#### 3.5.1 TonB1

Within the suite of genes from PA5531 – PA5541 discussed in sections 3.3 and 3.4, the increased abundance of PA5531, or TonB1, in Zn-depleted conditions, is of particular interest. TonB1 is typically considered to be associated with the uptake of iron-bound siderophores.

Though the closest same-strand regulator binding site upstream of the tonB1 gene is a ferric uptake regulator (Fur) box, the Zur binding site upstream of PA5536 is also located on this strand [25,89,90], and further, there does not appear to be a same-strand transcription terminator site between the Zur site and the tonB1 gene (Figure 6, as determined by analysis of genome features using Pseudomonas.com [25,91] and *P. aeruginosa* Zur-binding motifs [16]). Future work is needed to characterize the role PA5531 might play in Zn homeostasis, and/or the interplay of transcriptional regulators governing its expression.

#### 3.5.2 Cu homeostasis systems

A more well-characterized example of cross-metal homeostasis mechanisms is the intricate linkage of the *P. aeruginosa* Zn and Cu homeostasis systems (e.g. [92,93]). Several uptake and efflux mechanisms are known to transport both metals, and Zn resistance is reported to increase in response to Cu exposure [56,92–94]. Within this study, Cu chaperone CopZ2 (PA3574.1) increased in abundance under Zn exposure. CopZ2 binds Cu(II) ions with high-affinity in the cytoplasm and thus may function as a rapid response Cu-sequestering mechanism [95,96] and/or might facilitate the transfer of cytoplasmic Cu(II) to ATPases for efflux [93,97]. It is not clear what the benefit CopZ2 serves in conditions of Zn excess, but it may play a similar role through binding free Zn(II) ions to remove it from the cytoplasm. OpdT (PA2505, occD4), a putative tyrosine porin [98], also exhibited elevated abundance in Zn-replete samples. Consistent with our results, Teitzel et al. [98] and Wright et al. [100] reported Cu-dependent production of OpdT in *P. aerugionsa*, and Lee and colleagues [101] similarly noted an increase in its expression upon ZnO particle addition. The apparent paradoxical induction of a porin upon increased heavy metal exposure, as observed here, may be explained by its ability to perform low-level uptake of non-specific extracellular molecules: some authors [99,100] suggested that the decreased abundance of porins when faced with elevated metal concentrations is compensated for by the upregulation of OpdT.

#### 3.5.3 ColRS and ethanolamine metabolism

The ColRS TCS has recently been reported as responsive to Zn [102] in *Pseudomonas* species, and defects in this system result in disturbed proteomic profiles of Zn-exposed *Pseudomonas* cultures [76]. In line with these reports, upregulation of ColR occurred in Zn-replete conditions of this study with a 23-fold increase. In the presence of Zn, ColRS promotes remodeling of cell surface lipopolysaccharide component lipid A through addition of a phosphoethanolamine (pEtN) group, executed by pEtN transferase EptA [102]. Modification of the cell surface occurs in response to environmental stimuli, enabling a cell to adopt characteristics that grant it resistance to those stimuli. It is unclear how the modification of lipidA with pEtN promotes Zn resistance; it may simply be that other as-yet-unassessed genes controlled by the interaction of Zn with ColRS confer the desired resistance, and the stimulation of EptA is simply a byproduct [102]. However, a central ethanolamine catabolism system was also perturbed with variations in Zn concentration. HdhA (PA4022, exaC2), a hydrazone dehydrogenase, forms part of the PA4022-eat-eutBC operon, was enriched in Zn-replete samples. Together, these provide motivation to further investigate the role of Zn in modulating ethanolamine-based cell surface modifications.

## 4. Conclusions

Overall, this study provides further confirmation that revisiting biological systems using proteomic analyses can provide new insights and strengthen existing explanations of cellular behaviour. A global overview of the *P. aeruginosa* proteome when cultured in Zn-replete and Zn-depleted conditions revealed an apparent dichotomy in resource allocation between treatments: when Zn is available, a broader array of cellular functions is supported, whereas when Zn is depleted, the proteins produced in increased quantities are predominantly those involved in Zn trafficking. Through a combination of DDA and DIA analysis, we were able to detect many of the Zn homeostasis systems previously described. However, while we detected components of two suspected Zn ABC transporters, we did not detect any components of the ZnuABC transporter, suggesting that in *P. aeruginosa*, these other Zn uptake systems may play more dominant roles. Additionally, the sensitivity of these methods allowed us to observe the activation of metal homeostasis mechanisms not typically associated with Zn, which underscores the necessity of looking at cellular responses to stimuli holistically. Future investigations of metal allocation between proteins (such as through metalloproteomics techniques [2]) by *P. aeruginosa* under conditions of Zn excess and deprivation, and paired with knockout experiments of some of the genes discussed here, could shed light on the links between these systems.

## Supporting information

Systems Enrichment DIA

Systems Enrichment DDA

Processed Results

Genome Overview DIA

Metabolic Map DDA

Metabolic Map DIA

## Acknowledgements and Data Availability

The authors declare no conflict of interest. We thank the WHOI Ocean Ventures Fund and NIH Grant R01GM135709 for supporting this research. Thanks to the Manoil Lab for providing the *P. aeruginosa* culture. The mass spectrometry proteomics data have been deposited to the ProteomeXchange Consortium via the PRIDE partner repository with the dataset identifier PXD047454 and 10.6019/PXD047454.

## Supplemental

**Figure S1.**
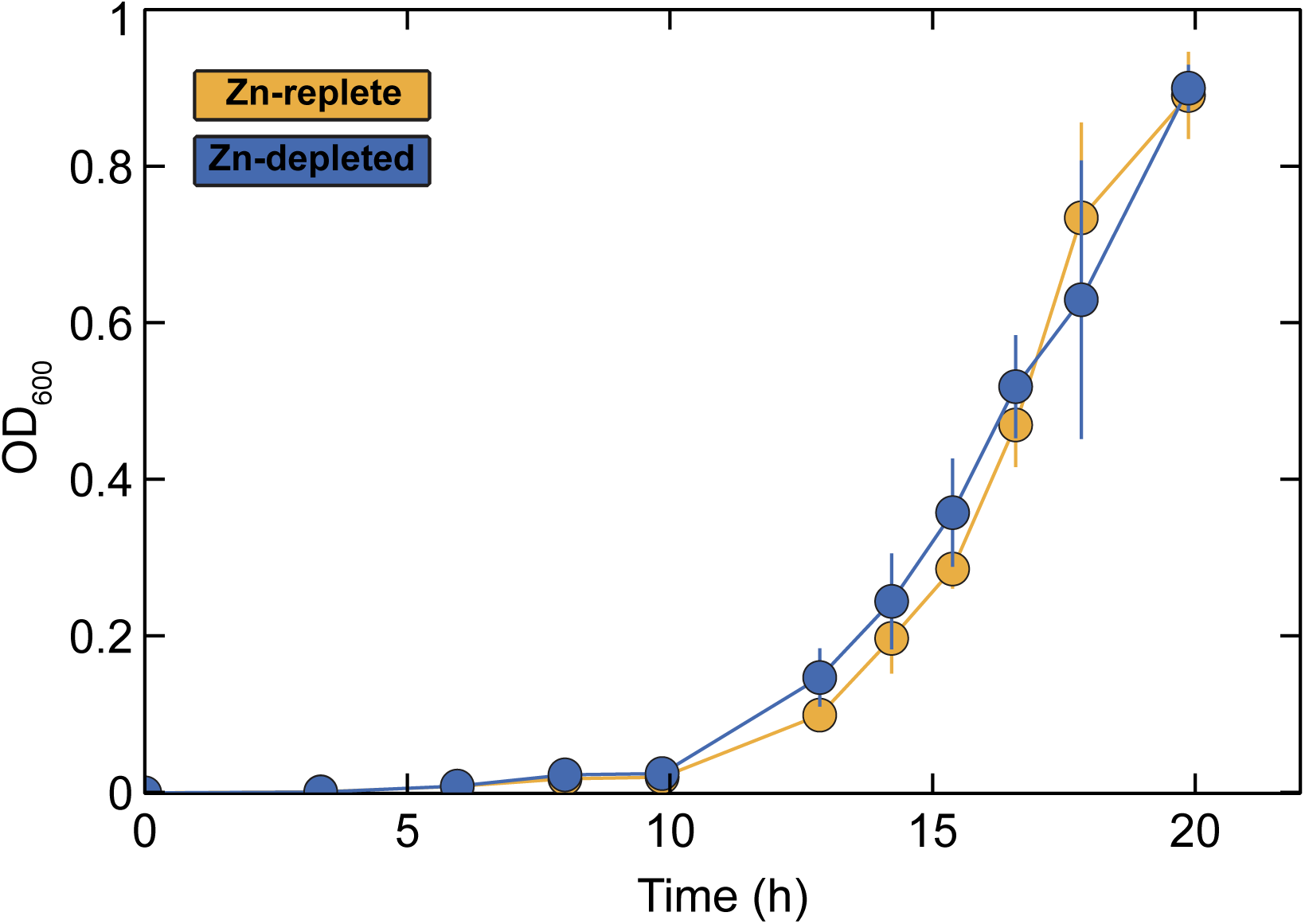
Growth curve of *P. aeruginosa* samples grown in Zn-replete (yellow) and Zn-depleted (blue) conditions, as determined by OD_600_ measurements. Error bars represent the standard deviation of biological triplicate samples.

## Alternate Text

Alt Text - Figure 1. Graphic showing the log_2_ fold-change values between Zn-depleted and Zn-replete samples. Values for each protein are indicated by colour scale and placed in the context of a graphical condensed representation of the *P. aeruginosa* PAO1 genome.

Alt Text - Figure 2. A modified bar graph shows the enrichment score of protein functional or pathway groups, with score indicated by line length, enrichment direction indicated by line type, false discovery rate indicated by colour, and gene count for each group indicated by marker size.

Alt Text – Figure 3. Bar graph with error bars and significance markers showing the abundance for various proteins involved in Zn acquisition for Zn-depleted and Zn-replete samples. For each protein, both DDA and DIA data are shown.

Alt Text – Figure 4. Bar graph with error bars and significance markers showing the abundance for various proteins involved in Zn efflux and tolerance for Zn-depleted and Zn-replete samples. For each protein, both DDA and DIA data are shown.

Alt Text – Figure 5. Bar graph with error bars and significance markers showing the abundance for Zn metalloproteins for Zn-depleted and Zn-replete samples, matched with the abundance of Zn-independent paralogs or functional analogs for both treatments.

Alt Text – Figure 6. Diagram illustrated the structure of *P. aeruginosa* PAO1 genome between PA5531 and PA5541. Direction of transcription, transcription factor binding sites and terminator sites are indicated. Color coding indicates whether the protein in question increased or decreased between treatments, or if it was undetected.

Alt Text – Supplemental Figure S1. Chart with error bars showing growth curve for *P. aeruginosa* grown in Zn-depleted and Zn-replete conditions.

Alt Text – Table 1. Table indicates the name and general function of proteins known to be involved in Zn uptake or efflux.

Alt Text – Table 2. Table shows the log_2_ fold-change and associated statistical information for proteins discussed in this paper as measured by DDA and DIA methods.

