## Supplementary material for "Proteomic profiling of zinc homeostasis mechanisms in *Pseudomonas aeruginosa* through data-dependent and data-independent acquisition mass spectrometry": Genome Overview DIA

### *Pseudomonas aeruginosa* PAO1 Genome Overview

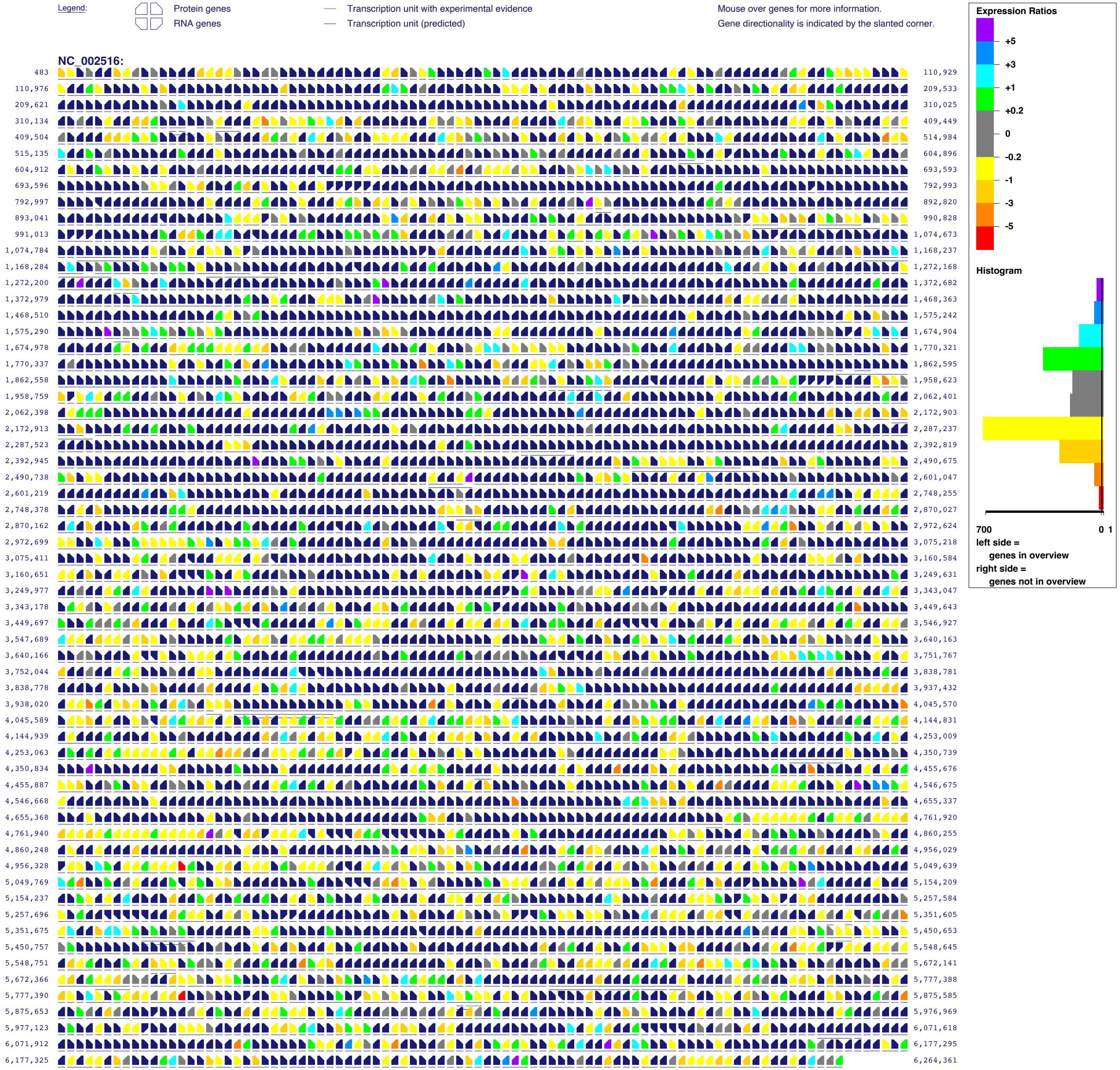


Generated by SRI International Pathway Tools version 28.0 on Tue Dec 3, 2024
