## Supplementary figures and images for "Proteomic profiling of zinc homeostasis mechanisms in *Pseudomonas aeruginosa* through data-dependent and data-independent acquisition mass spectrometry"

### Metabolic Map DDA

***Pseudomonas aeruginosa* PAO1 Cellular Overview**

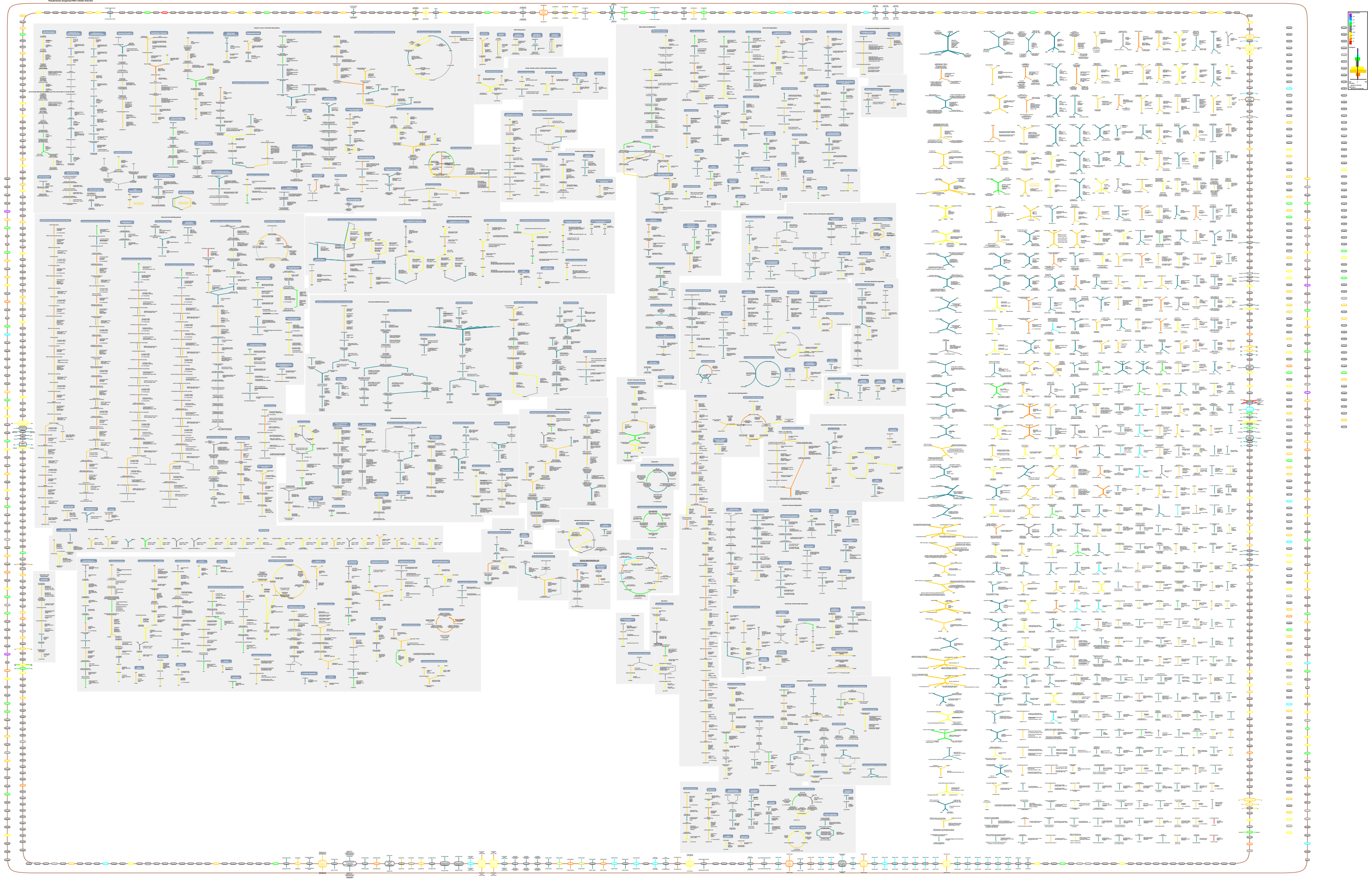

### Metabolic Map DIA

*Pseudomonas aeruginosa* PAO1 Cellular Overview

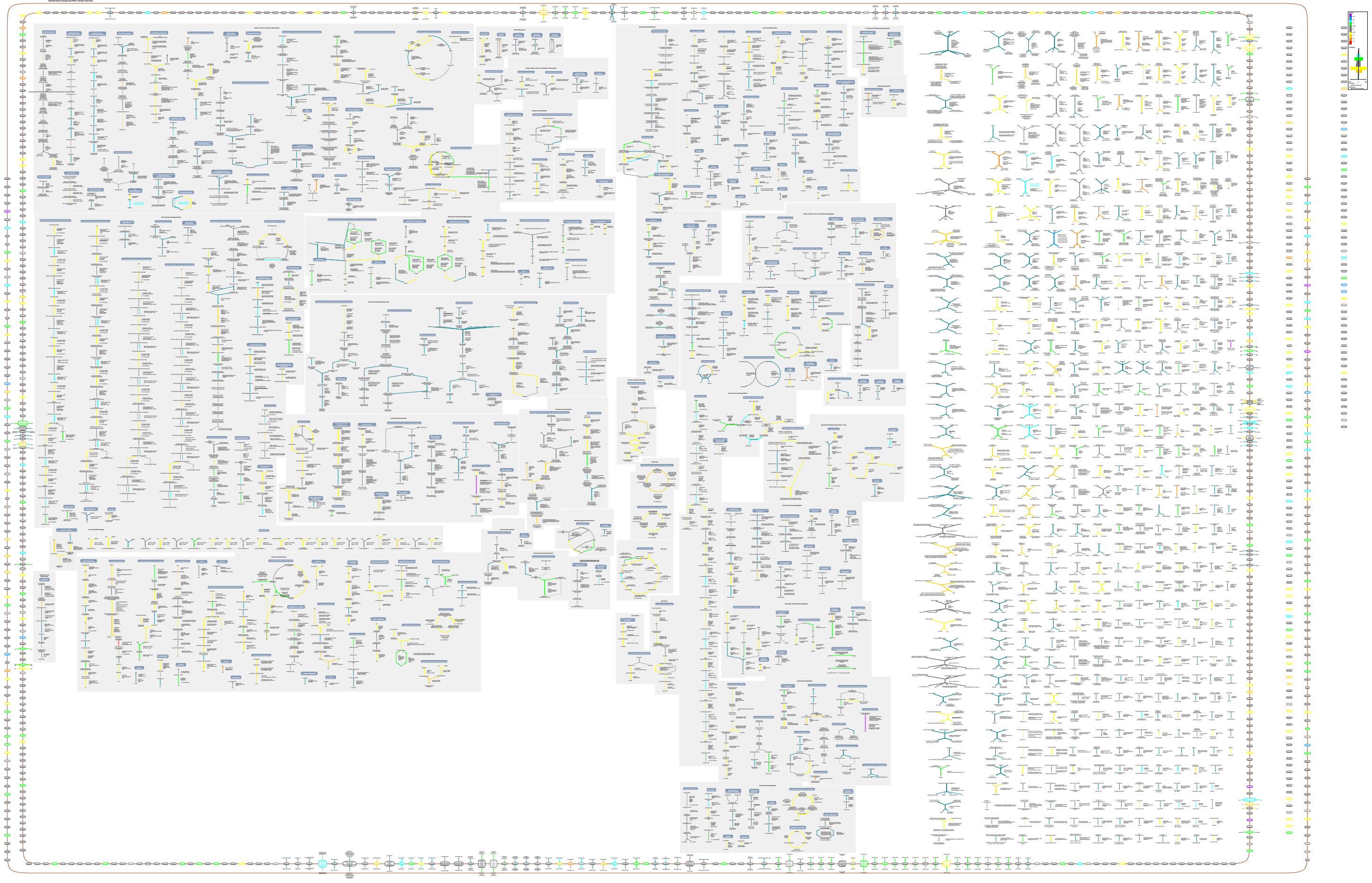
